# Genetic divergence and phenotypic plasticity contribute to variation in cuticular hydrocarbons in the seaweed fly *Coelopa frigida*

**DOI:** 10.1101/303206

**Authors:** Emma Berdan, Swantje Enge, Göran M. Nylund, Maren Wellenreuther, Gerrit A. Martens, Henrik Pavia

## Abstract

Cuticular hydrocarbons (CHCs) form the boundary between insects and their environments and often act as essential cues for species, mate and kin recognition. This complex polygenic trait can be highly variable both among and within species, but the causes of this variation, especially the genetic basis, are largely unknown. In this study, we investigated phenotypic and genetic variation of CHCs in the seaweed fly, *C. frigida*, and found that composition was affected by both genetic (sex and population) and environmental (larval diet) factors. We subsequently conducted behavioral trials that show CHCs are likely used as a sexual signal. We identified general shifts in CHC chemistry as well as individual compounds and found that the methylated compounds, mean chain length, proportion of alkenes, and normalized total CHCs differed between sexes and populations. We combined this data with whole genome re-sequencing data to examine the genetic underpinnings of these differences. We identified 11 genes related to CHC synthesis and found population level outlier SNPs in 5 that are concordant with phenotypic differences. Together these results reveal that the CHC composition of *C. frigida* is dynamic, strongly affected by the larval environment, and likely under natural and sexual selection.

## INTRODUCTION

In insects, cuticular hydrocarbons (CHCs) are a primary adaptation to life on land because they protect against desiccation (Wigglesworth 1945; Blomquist and Bagnères 2010). However, in many solitary and social insects, cuticular hydrocarbons are also used as one of the primary cues to recognize, and possibly discriminate between species, sexes, and among kin (Bagneres et al. 1996; Ferveur 2005; Blomquist and Bagnères 2010; Van Oystaeyen et al. 2014). The multifarious use of CHCs in adaptation and communication means that the composition is frequently under both natural (Howard and Blomquist 2005; Blomquist and Bagnères 2010; Foley and Telonis-Scott 2011; Rajpurohit et al. 2017) and sexual selection (Ferveur 2005; Howard and Blomquist 2005; Peterson et al. 2007; Thomas and Simmons 2009; Blomquist and Bagnères 2010; Steiger et al. 2013).

*De novo* synthesis of CHCs is well defined and conserved across insects (Howard and Blomquist 2005; Blomquist and Bagnères 2010): Synthesis takes place in oenocytes and starts with acetyl-CoA which is then elongated by a fatty acid synthase (FAS) forming a long-chained fatty acyl-CoA. FASs may also add methyl groups to these fatty acids (de Renobales et al. 1986; Blomquist et al. 1994; Juarez et al. 1996; Chung et al. 2014). Elongases further lengthen these fatty acyl-CoAs and double bonds or triple bonds are added by desaturases (Howard and Blomquist 2005; Blomquist and Bagnères 2010). These are reduced to aldehydes by reductases and then these aldehydes are converted to hydrocarbons by a decarboxylation reaction, which is mediated by a cytochrome P450 (Qiu et al. 2012). Coding or expression variation in any number of these enzymes can lead to significant changes in CHC composition (ex: de Renobales et al. 1986; Reed et al. 1995; Qiu et al. 2012; Chung et al. 2014; Dembeck et al. 2015; Chen et al. 2016).

Changes in gene expression (Dallerac et al. 2000a; Chertemps et al. 2007; Shirangi et al. 2009; Feldmeyer et al. 2014) and/or coding sequences (Keays et al. 2011; Badouin et al. 2013; Kulmuni et al. 2013) can impact both the overall composition of CHCs and the amount of specific compounds. Multiple environmental factors may also influence CHC composition. For example, CHCs can vary under different temperature regimes (Toolson and Kupersimbron 1989; Savarit and Ferveur 2002; Rouault et al. 2004), diets (Stennett and Etges 1997; Liang and Silverman 2000; Stojkovic et al. 2014) or other climatic variables (Howard et al. 1995; Kwan and Rundle 2010). Moreover, multiple studies have found that CHC composition varies between natural populations (ex: Haverty et al. 1990; Etges and Ahrens 2001; Frentiu and Chenoweth 2010; Perdereau et al. 2010), but it is unknown how much of this variation is due to plasticity versus genetic divergence. The contribution of different factors can be teased apart using laboratory studies. The role of genetic factors can, for example, be assessed by raising different populations in the same environment. Likewise, plasticity can be assessed by raising the same population in multiple environments (i.e. common garden and reciprocal transplant experiments). Determining the causes and consequences of CHC variation is an important general step towards understanding the evolution of CHCs and their co-option as sexual signals.

The seaweed fly *Coelopa frigida* presents an attractive system in which to investigate causes and consequences of variation in CHCs. These flies live in highly dynamic environments (Berdan et al. 2018) making it likely that both genetic changes as well as phenotypic plasticity affect CHC composition. *Coelopa frigida* lives in “wrackbeds”, accumulations of decomposing seaweed on shorelines that act as both habitat and food source for larvae and adults. Local adaptation in response to wrackbed composition has been demonstrated in Swedish *C. frigida* (Wellenreuther et al. 2017). Furthermore, CHCs may be under sexual selection as it is possible that CHCs are used as a mating signal in this system. Mating in *C. frigida* is characterized by intense sexual conflict with no courtship behaviour but evidence of female choice. Males will approach and forcefully mount females. Females are reluctant to mate and use a variety of responses to dislodge the male, such as downward curling of the abdomen (to prevent contact), kicking of the legs, and shaking from side to side (Day et al. 1990; Blyth and Gilburn 2011). It has been consistently reported that mating attempts by large males are more likely to result in copulation (Butlin et al. 1982; Gilburn et al. 1992, 1993) but the signals used for mate choice are unknown.

Here we investigate the cuticular hydrocarbons of *C. frigida.* Specifically, we examine variation in CHC signatures due to sex (possible signature of sexual selection), population (possible signature of natural selection), and larval environment/diet (phenotypic plasticity). We combine this with behavioural trials to test the significance of CHCs in a mate choice context. We also investigate genetic variation that may underlie this phenotypic variation. We discuss our findings in light of sexual selection and adaptation in this species and others and outline future research areas that deserve attention.

## METHODS

### Study Species

*Coelopa frigida* belongs to the group of acalyptrate flies which exclusively forage on decomposing seaweed (Cullen et al. 1987). This species is found along the seashores of Northern Europe (Mcalpine 1991), plays a vital role in coastal environmental biodiversity (Griffin et al. 2018) and health by accelerating the decomposition of algae, allowing for faster release of nutrients (Cullen et al. 1987).

### Sampling

Fly larvae were collected in April and May 2017 from two Norwegian populations, Skeie (58.69733, 5.54083) and Østhassel (58.07068, 6.64346), and two Swedish populations, Stavder (57.28153, 12.13746) and Ystad (55.425, 13.77254). These populations are situated along an environmental cline in salinity, wrackbed composition, and wrackbed microbiome (Berdan et al., in prep; Day et al. 1983) from the North Sea to the Baltic Sea (Figure 1). Larvae were transported to Tjärnö Marine Laboratory of the University of Gothenburg where they completed development in an aerated plastic pot filled with their own field wrack in a temperature controlled room at 25 °C with a 12h/12h light-dark cycle. After adults eclosed in the lab, they were transferred to a new pot filled with standard wrack consisting of 50 % *Fucus spp*. and 50 % *Saccharina latissima* which had been chopped, frozen, and then defrosted. The exception to this is a subset of Ystad adults that were allowed to mate on the same material they had been collected in (field wrack). Thus we had 5 treatment groups: All four populations (Skeie, Østhassel, Stavder, Ystad) on standard wrack, and Ystad also on field wrack. Adult flies were allowed to lay eggs on the provided wrack and this second generation was raised entirely in a temperature controlled room at 25 °C with a 12 h/12 h light dark cycle. As larvae pupated they were transferred to individual 2 mL tubes with a small amount of cotton soaked in 5 % (w/v) glucose to provide moisture and food for the eclosing adult. This ensured virginity in all eclosing flies. The tubes were checked every day and the date of eclosure of each fly was noted. Two days after eclosure adult flies were frozen at −80 °C and kept there until CHC extraction.

**Figure 1.**
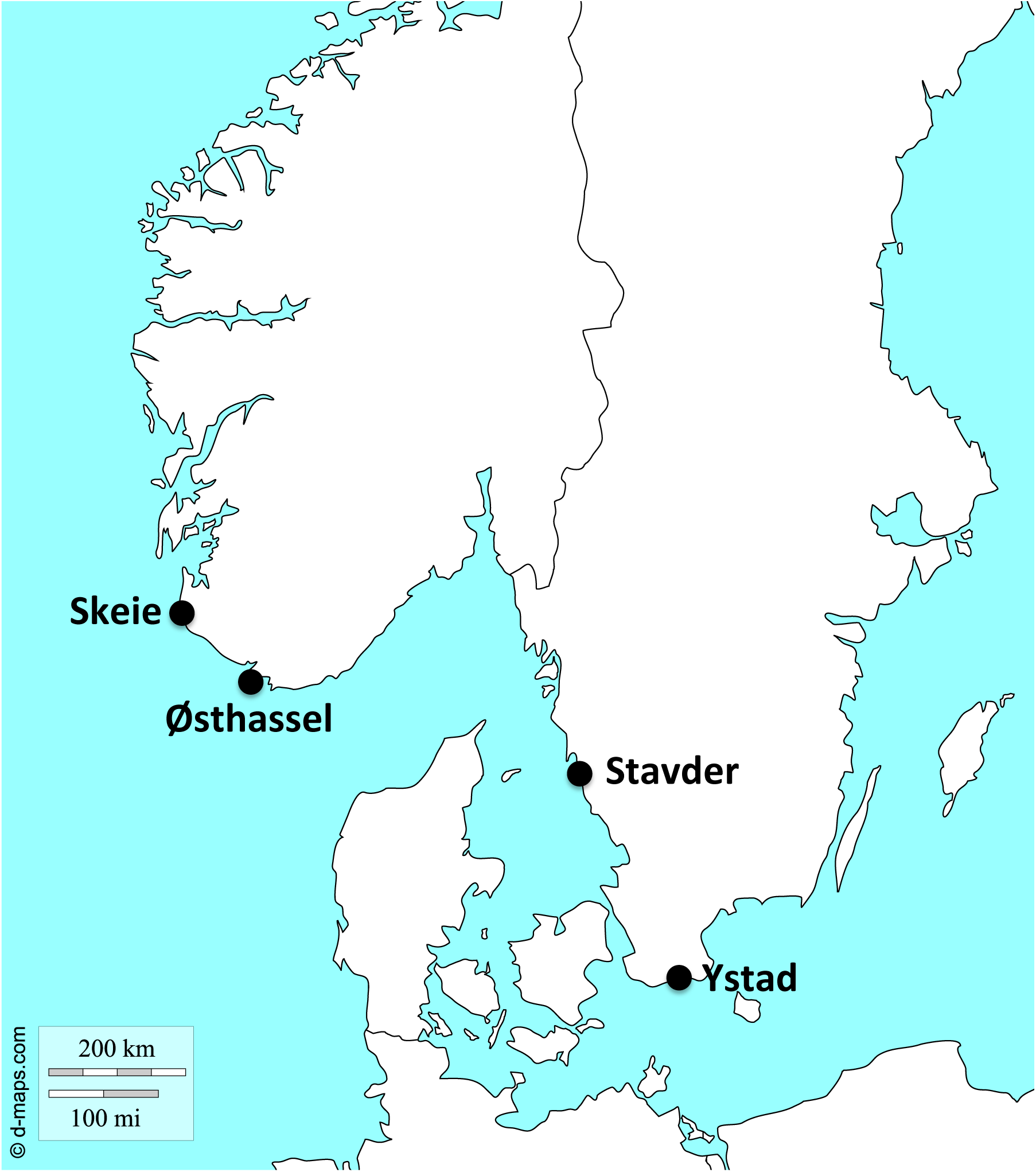
Map of *C. frigida* populations used in this study. Base map from d-maps (http://d-maps.com/carte.php?num_car=5972)

### CHC Analysis

Frozen flies were placed in 12-well porcelain plates to defrost and dry, as moisture can affect lipid dissolution. Each fly was then placed in a 1.5 ml high recovery vial containing 300 µl of *n*-hexane, vortexed at a low speed for 5 second, and extracted for 5 minutes. Afterwards flies were removed from the vial and allowed to air dry on the porcelain plates before they were weighed (Sartorius Quintix 124-1S microbalance) to the nearest 0.0001 grams and sexed. Extracts were evaporated until dry under nitrogen gas and then stored at −20°. Before analysis 20 µl of *n*-hexane containing 1 µg/ml *n*-nonane was added as an internal standard to each vial, which was then vortexed at maximum speed for 10 seconds.

The extracts from 20 male and 20 female flies from every treatment group were analyzed on a GC (Agilent GC 6890) coupled to a MS (Agilent 5973 MSD) using a HP-5MS capillary column (Agilent) (see Table S1 in supplementary materials for instrument settings).

The total ion chromatograms were quality checked, de-noised and Savitzky-Golay filtered before peak detection and peak area integration in OpenChrom®. Peaks were then aligned using the R-package ‘GCalignR’(Ottensmann et al. 2018) prior to statistical analysis. Peaks were tentatively identified by their mass spectra and retention time index based on a C21-C40 *n-*alkane standard solution (Sigma-Aldrich).

### Statistical Analysis

The effects of sex and population on CHC profile were assessed using flies raised on standard wrack using a balanced sampling design (N=13 for each combination) and 127 peaks were identified by the R package “GCalignR”. The effects of sex and diet (i.e. wrack type) during the larval stage on CHC profiles were analyzed using a balanced design of flies from the Ystad population (N=12 for each combination) and 111 peaks were identified by R package “GCalignR”. Peak areas were normalized on the peak area of the internal standard and the weight of the fly before being auto-scaled prior to statistical analysis. Clustering of samples was visualized with a PCA, and group differences were analyzed using a PERMANOVA followed by multiple group comparisons using the PRIMER-E 7 software. We also analyzed group differences via (O)PLS-DA using the “ropls”-packages in R (Thevenot et al. 2015). Candidate compounds for group differentiation were determined from variables of importance for OPLS-DA projection (VIP scores) (Galindo-Prieto et al. 2014), which were further assessed by univariate analysis using a false discovery adjusted significance level of α = 0.05 using the Benjamini and Hochberg’s FDR-controlling procedure (Benjamini & Hochberg, 1995).

We examined sex and country effects in more depth to examine shifts in chemistry that could be linked to our genetic data (see below). Our results above indicated that the differences in CHC profiles between sexes and populations of *C. frigida* were due to quantitative rather than qualitative differences (i.e. changes in relative amounts rather than presence/absence of certain compounds). Thus, we looked for general shifts in CHC chemistry by examining 1. The total normalized peak area (i.e. the sum of all peaks used in analysis), 2. The proportion of alkenes, and 3. The proportion of methylated compounds. For #1 we used all 127 peaks used in the previous analysis and for #2 and #3 we used the subset of these peaks that could be accurately categorized (116 peaks). For the total peak area we transformed the data with a log transformation and then used the ‘dredge’ function from the MuMIn package (Barton 2017) to compare AIC values for nested glm (Gaussian distribution, link=identity) models with the terms sex, country, and country*sex. If there were two models that differed in AIC by less than 1, we chose the simpler model. For alkenes and methylated compounds our data was proportional and did not include 1 or 0 so we used the ‘betareg’ function from the betareg package (Grun et al. 2012). We again used the ‘dredge’ function to compare AIC values for nested models with the terms sex, country, and country*sex and took the simpler model in case of an AIC difference of less than 1.

We also analysed the distribution of chain lengths in our samples. To do this, we subsetted all peaks where we had accurate length information (107 peaks) and then calculated the weighted mean chain length as

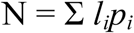

where *l_i_* is the length of a carbon chain and *p_i_* is the proportional concentration of all chains of that length. We also calculated dispersion around this mean as

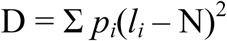

We analyzed these in the same manner as described above, using AIC values to conduct model choice from glm models (Gaussian distribution, link=identity).

### Behavioral Trials

Given experimental restrictions, we were unable to accurately test for the use of CHCs in female choice and thus we tested for the use CHCs in male choice. Behavioral trials were conducted with flies from the Østhassel population that were collected in March 2018 and cultured in the lab for several generations. To collect adults for testing, we selected pupae and kept them in individual Eppendorf tubes (to ensure virginity) containing a small piece of cotton soaked in a 0.5% mannitol solution. When a fly eclosed, the Eppendorf was moved to the refrigerator to slow down aging. In all behavioural trials, five males (marked individually with nail polish) were introduced to a small petri dish (Ø 55mm) with a small amount of minced *Saccharina* (≈1g) and two dead flies mounted on insect pins. We conducted two different types of trials: I. Males were able to choose between a dead male and a dead female and II. Males were able to choose between two dead females one of which had been extracted in hexane (see above for methods). Type 1 trials were used to confirm that our methodology was working and type II trials were used to test if CHCs play a role in male choice. For type II trials we checked that extracted females were mostly devoid of CHCs by re-extracting five females and performing GC-MS analysis as described above. The hexane extraction reduced the total amount of CHCs on the flies by 72% ±6. All trials lasted for 30 minutes and we recorded all mountings and how long they lasted. We used a paired Wilcoxon rank-sum test on our data, with male as our unit of replication, to determine if males preferentially mounted one of the two proffered flies.

### Candidate gene analysis

Based on a literature search for genes involved in CHC synthesis, we identified 37 reference protein sequences, which were downloaded from Flybase (flybase.org) in October 2017. We used tblastx with the default settings to identify orthologs in the *C. frigida* transcriptome (Berdan & Wellenreuther, unpublished). We then used gmap (Wu and Watanabe 2005) to find areas of the *C. frigida* genome (Wellenreuther, *et al*., unpublished) that might contain these genes. We output all alignments as a gff3 file. We have a previously made a VCF file created from 30x whole genome re-sequencing of 46 *C. frigida* across five populations in Scandinavia (Berdan *et. al*, unpublished). This VCF file was made by aligning raw reads to the *C. frigida* genome using bwa-mem (Li and Durbin 2010), duplicate reads were marked using ‘picard’ (http://picard.sourceforge.net). SNPs were called with the Genome Analysis Toolkit (GATK; DePristo *et al*. 2011; Van der Auwera *et al*. 2013) specifically the GATK-module ‘UnifiedGenotyper’ (Van der Auwera *et al*. 2013). VCF filtering was done as described in Berdan *et al*. 2015. We used this VCF file and the gff3 output from gmap in SNPeff (Cingolani et al. 2012) to annotate the SNPs in these genes.

To find which SNPs are diverged between populations we used vcftools (Danecek et al. 2011) to calculate Cockerham and Weir’s F_ST_ estimate (Weir and Cockerham 1984) for all SNPs in our reference genome. We retained the top 5% of this distribution (F_ST_ ≥ 0.142, mean F_ST_ 0.024) as SNPs that were potentially divergent. We then subset the VCF file to include only outlier SNPs located in our previously identified genomic regions.

## RESULTS

### CHC analysis

A diverse blend of hydrocarbons with more than 100 different compounds was found in the cuticular extract of *C. frigida*. The majority of CHCs ranged in chain length from 23 to 33 carbon atoms. They contained odd numbered *n-*alkanes, methyl-, dimethyl- and trimethyl-alkanes, as well as odd numbered alkenes. Some even numbered alkanes were also present but in lower quantities.

### Sex and population effects

Results of the PERMANOVA indicated significant differences between females and males and between populations but also a significant interaction between sex and population (Table 1A). Pairwise tests on the interaction showed a significant difference between males and females for all populations except Ystad (Table S2). Multiple comparisons of populations showed that the Norwegian populations (Skeie and Østhassel) were similar while the Swedish populations (Ystad and Stavder) differed for males but not for females. In general the Norwegian populations tended to differ from the Swedish and a similar pattern was also shown in the results of the PCA and PLS-DA (Figure 2A, C, Table S3). After combining the Swedish and Norwegian populations, the PERMANOVA indicated highly significant differences between females and males as well as between Sweden and Norway (Table 1B). OPLS-DA models for both sex and country were significant and lead to separate clusters of females and males, and Sweden and Norway, respectively (Figure 2B, D, Table S3). The OPLS-DA model revealed 41 peaks that contributed with a VIP score >1 to the differentiation between females and males. Of these, 31 were also significant in t-tests with a FDR adjusted p-values (Table 2). Generally compounds were more abundant in males with the exception of only 3 peaks that were more abundant in females. Looking at country differentiation between Swedish and Norwegian populations we found 43 compounds with a VIP score >1, of which 29 were significant in t-tests with FDR adjusted p-values and consisted of an equal mix of compounds more abundant in either of the countries (Table 2). Differences in CHC profiles between sexes and populations were due to quantitative differences, affecting a wide range of compounds instead of qualitative differences, i.e. the presence/ absence of a few distinct compounds

**Table 1.** PERMANOVA and PERMDISP results for A. Sex and Population, B. Sex and Country, and C. Sex and Wrack type. * < 0.05 ** < 0.01 *** < 0.001

| A. PERMANOVA: Sex and Population |  |  |  |  |  | PERMDISP |
| --- | --- | --- | --- | --- | --- | --- |
|  | df | SS | MS | F | p-value |  |
| Sex | 1 | 658.1 | 658.1 | 5.86 | <0.001 *** | see S2 |
| Population | 3 | 1150.3 | 383.4 | 3.41 | <0.001 *** |  |
| Sex:Population | 3 | 486.9 | 162.3 | 1.44 | 0.015 * |  |
| Residuals | 96 | 10786.0 | 112.4 |  |  |  |
| Total | 103 | 13081 |  |  |  |  |

| B. PERMANOVA: Sex and Country |  |  |  |  |  | PERMDISP |  |
| --- | --- | --- | --- | --- | --- | --- | --- |
|  | df | SS | MS | F | p-value | F | p-value |
| Sex | 1 | 658.0 | 658.1 | 5.60 | <0.001 *** | 8.94 | 0.003 ** |
| Country | 1 | 517.8 | 517.8 | 4.41 | <0.001 *** | 1.19 | 0.279 |
| Sex:Country | 1 | 154.1 | 154.1 | 1.31 | 0.128 |  |  |
| Residuals | 100 | 11751.0 | 117.5 |  |  |  |  |
| Total | 103 | 13081 |  |  |  |  |  |

| C. PERMANOVA: Sex and Wrack type |  |  |  |  |  | PERMDISP |  |
| --- | --- | --- | --- | --- | --- | --- | --- |
|  | df | SS | MS | F | p-value | F | p-value |
| Sex | 1 | 360.9 | 360.9 | 3.73 | <0.001 *** | 2.36 | 0.129 |
| Wrack | 1 | 481.3 | 481.3 | 4.98 | <0.001 *** | 2.29 | 0.139 |
| Sex:Wrack | 1 | 122.3 | 122.3 | 1.27 | 0.1629 |  |  |
| Residuals | 44 | 4252.5 | 96.7 |  |  |  |  |
| Total | 47 | 5217 |  |  |  |  |  |

**Figure 2.**
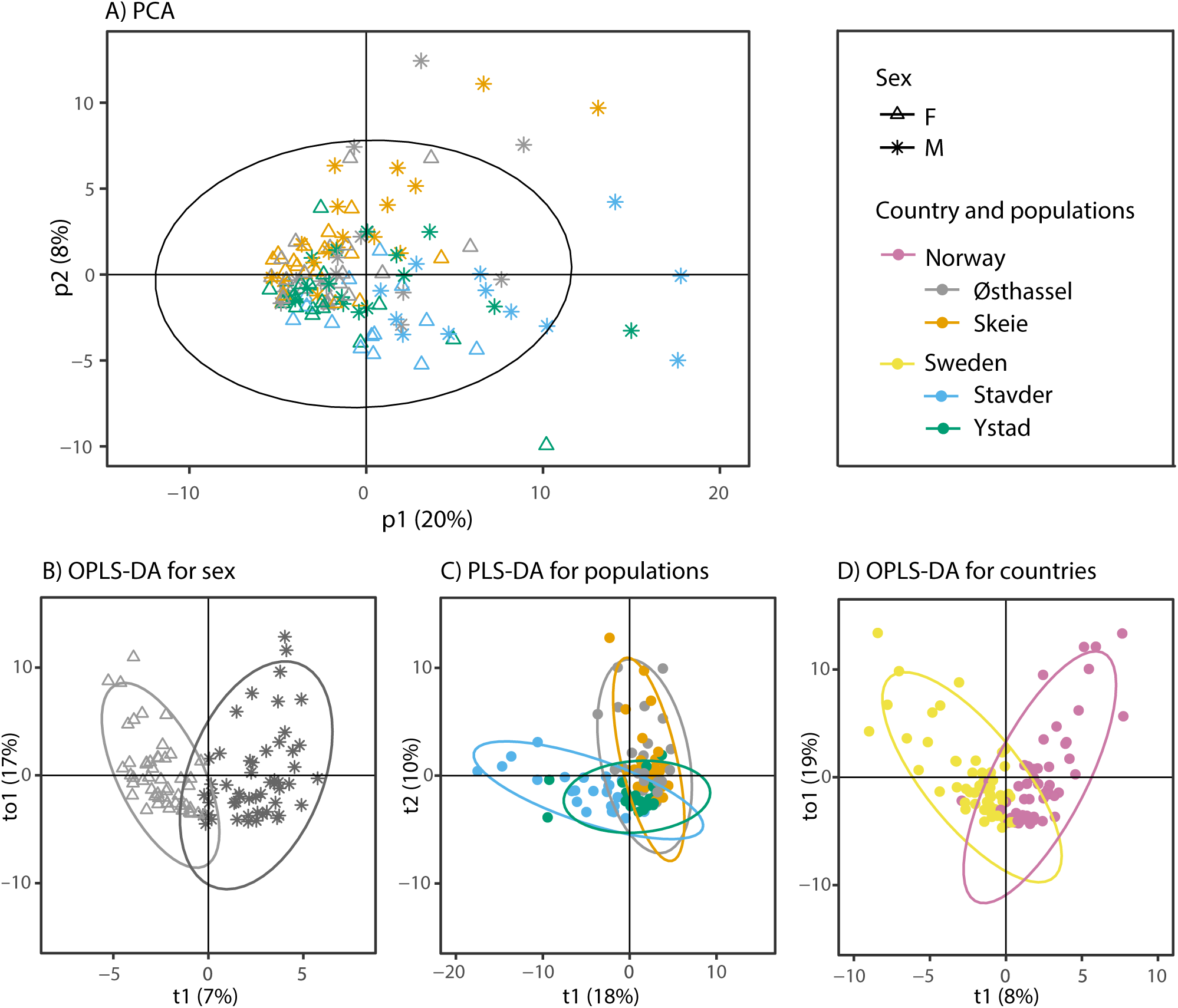
A. PCA of all samples from the balanced data set indicating population (grey – Østhassel, orange – Skeie, blue – Stavder, green – Ystad) and sex (triangle – female, star – male). B. OPLS-DA for sex using all samples from the balanced data set, C. PLS-DA for population using all samples from the balanced data set, and D. OPLS-DA for country using all samples from the balanced data set (yellow – Sweden, purple – Norway). Circles indicate 95% confidence regions.

**Table 2.**
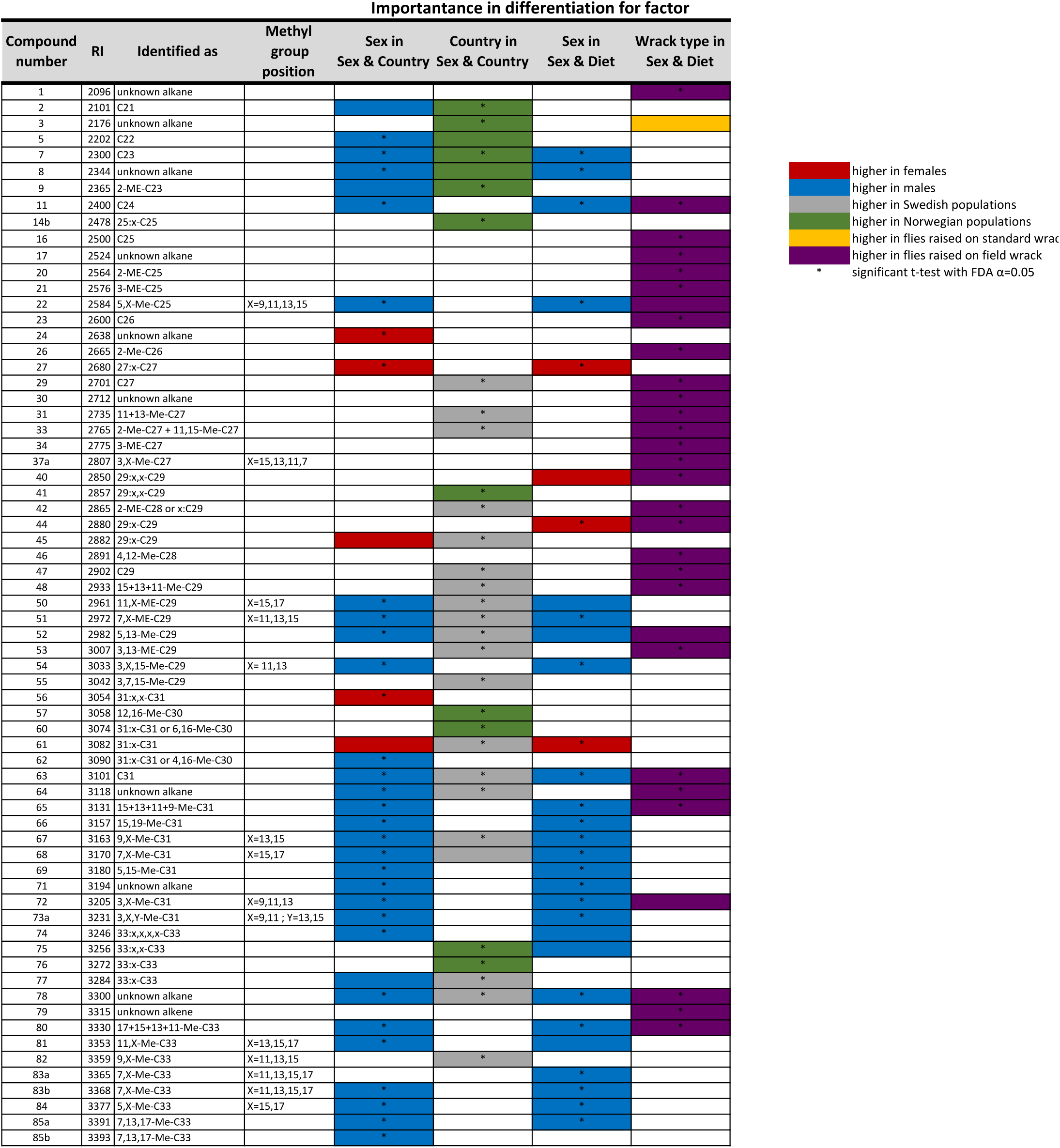
List of compounds that are significantly important for differentiation of at least factor (sex, diet, or country). If compounds are significant for differentiation of any category in any comparison a * appears under that factor. Color indicates the direction of the differentiation (red-higher in females, blue – higher in males, grey – higher in Norwegian populations, green – higher in Swedish populations, yellow – higher in flies raised on standard wrack, purple – higher in flies raised on field wrack).

### Sex and diet effects

PERMANOVA demonstrated a significant difference between females and males in the Ystad population and also a significant effect of the larval diet (wrack type) (Table 1C). Both OPLS-DA models for either sex or wrack type were significant and lead to separate clusters of females and males and flies raised on standard or field wrack (Figure 3B,C, Table S3). The OPLS-DA model revealed 38 peaks that contributed to the differentiation between females and males with a VIP score >1. Of these, 24 compounds were also significant in a t-test with FDR adjusted p-values (Table 2). All of these, except for 3 peaks, were more abundant in males. Overall 65% (26 compounds) were shared between this analysis and the previous analysis investigating sex and population effects. Looking at differentiation between flies raised on a diet consisting of standard or field wrack we found 33 compounds with a VIP score >1. Of these, 27 compounds were significant using a t-test with FDR adjusted p-values, all with higher amounts in flies reared on field wrack (Table 2). Flies reared on field wrack had more CHCs per gram body weight than flies reared on standard wrack (normalized totals: Females field wrack 7.96 ± 0.94, females standard wrack 4.09 ± 0.47, males field wrack 11.24 ± 1.03, males standard wrack 7.09 ± 1.21). Differences in CHC profiles between sexes and diets were exclusively due to quantitative differences, affecting a wide range of compounds instead of qualitative differences, i.e. the presence/ absence of a few distinct compounds

**Figure 3.**
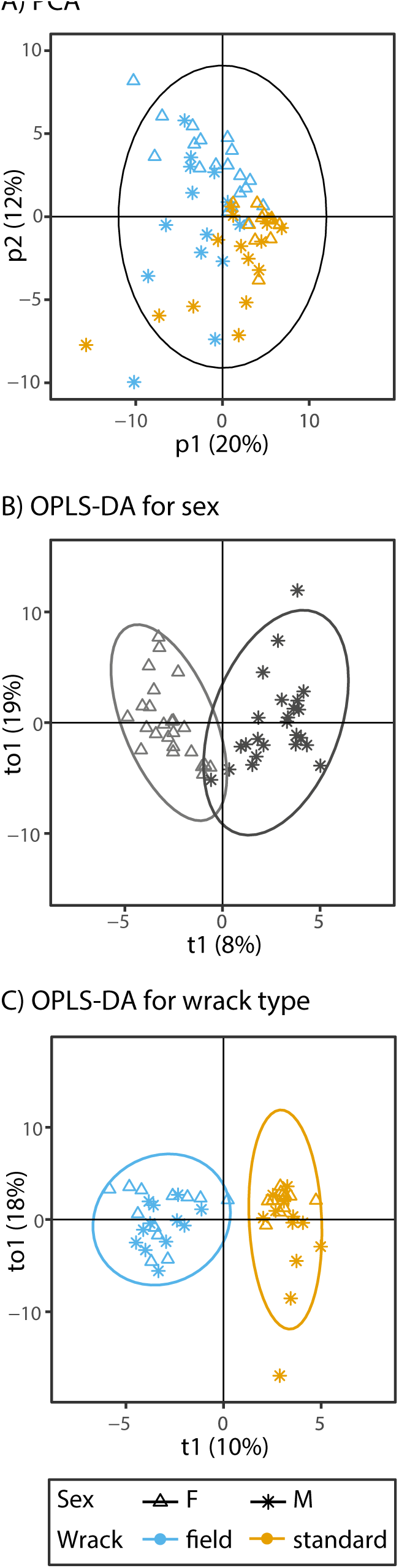
A. PCA of samples from Ystad indicating diet (blue – field, yellow – standard) and sex (triangle – female, star – male). B. OPLS-DA for sex using only Ystad samples. C. OPLS-DA for diet using only Ystad samples. Circles indicate 95% confidence regions.

### Shifts in chemistry

As our results above indicated an overall effect of country rather than population we used country and sex for our terms in the glm models. For total normalized peak area, the best model contained only sex as a main effect (for a full AIC comparison for all models see Table S4A-E). In general, males had more CHCs per gram body weight than females (Figure 4A). For the proportion of alkenes, the best model contained country and sex as main effects (for type II analysis of deviance tables for all best models see Table S5A-E). Females tended to have more alkenes than males and individuals from Norway had more alkenes than individuals from Sweden (Figure 4B). The best model for the proportion of methylated compounds contained sex and country as the main effects. As with alkenes, females tended to have more methylated compounds than males and individuals from Norway tended to have more methylated compounds than individuals from Sweden (Figure 4C).

**Figure 4.**
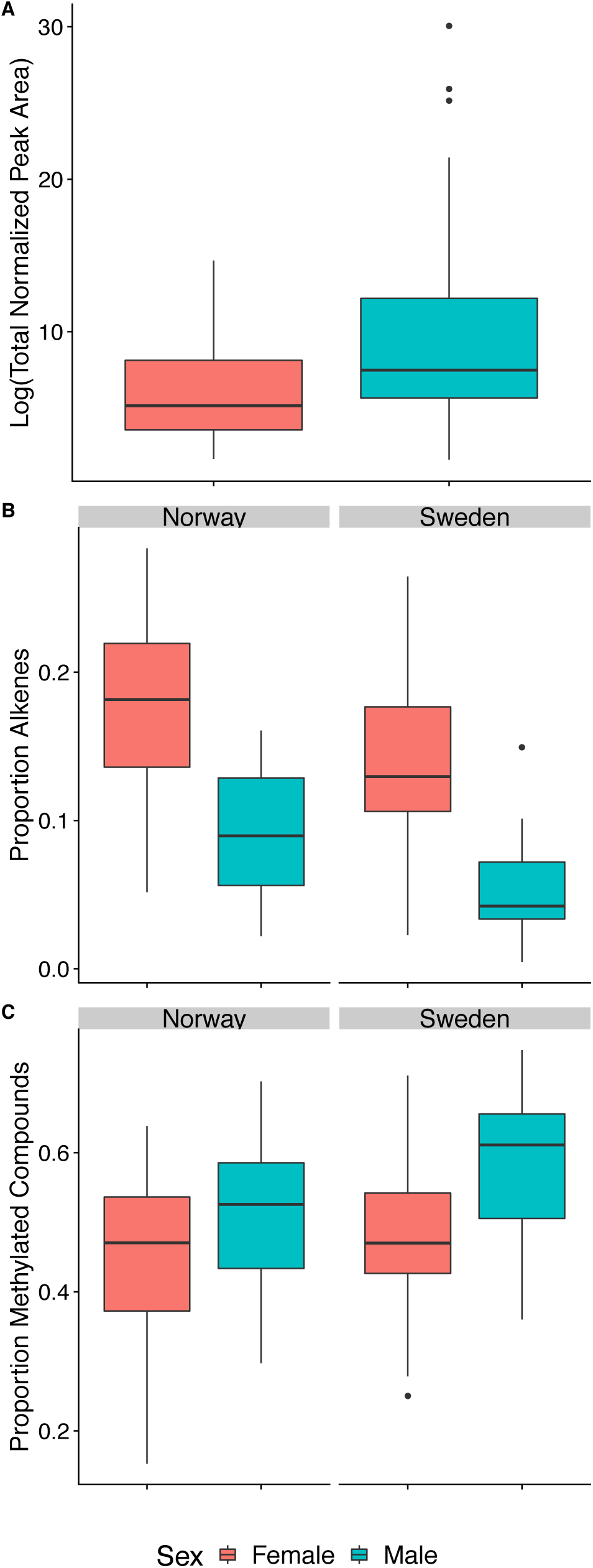
Boxplot showing differences in A. Log transformed total peak area, B. proportion of compounds that are alkenes, C. proportion of methylated compounds. Females are in red, males are in blue.

The best model for weighted mean chain length was country and sex. Overall, males had longer mean chain lengths than females and individuals from Sweden had longer mean chain lengths than individuals from Norway (Figure 5A). The best model for dispersion around mean chain length was also country and sex. Males had higher dispersion than females and individuals from Norway had higher dispersion than individuals from Sweden (Figure 5B).

**Figure 5.**
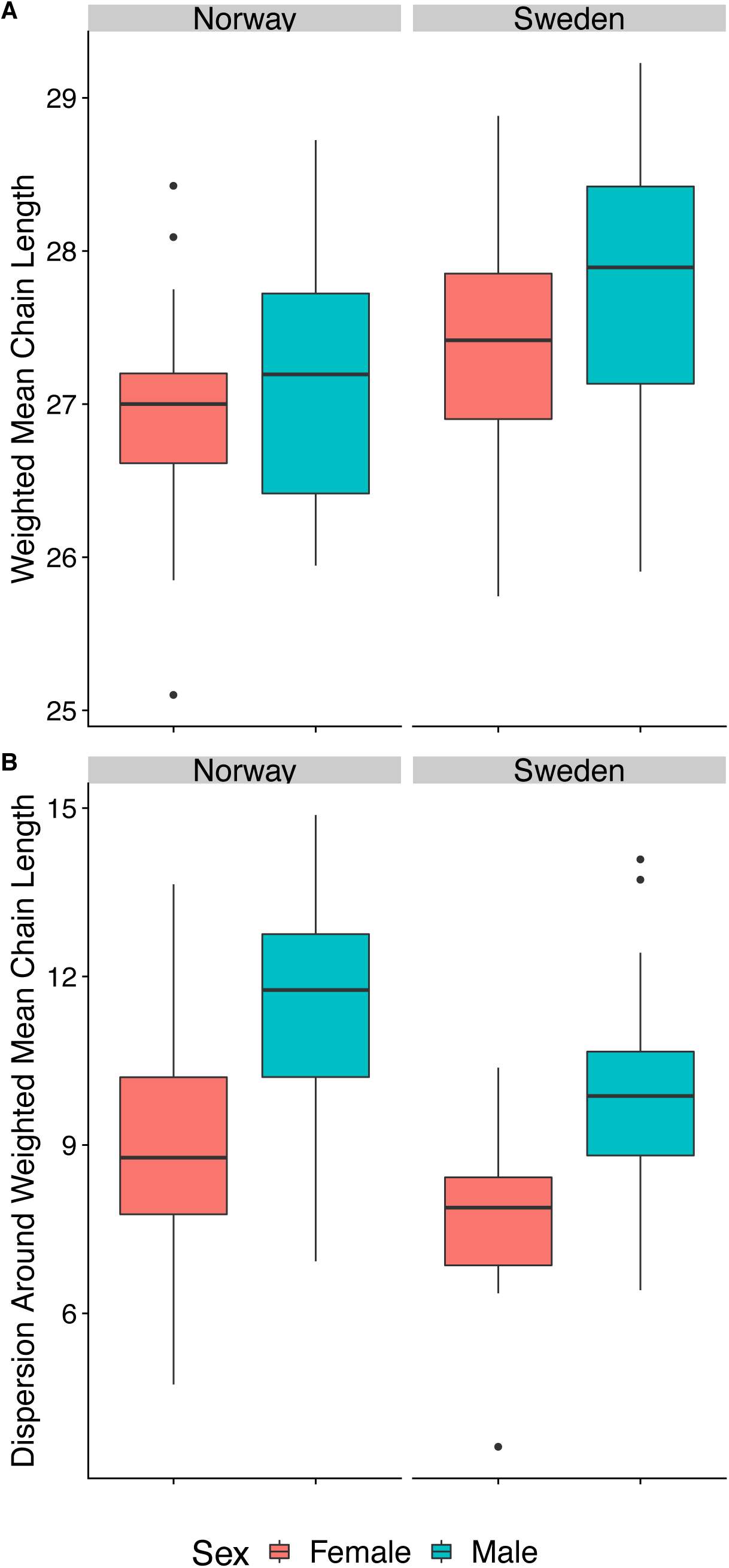
Boxplot showing differences in A. Weighted mean chain length and B. Dispersion around the mean. Females are in red, males are in blue.

### Behavioural trials

We conducted 20 trials of type I (female vs. male) and 12 trials of type II (female vs. extracted female). We only retained data from males that mounted at least one of the proffered flies leaving us with data from 28 males (type I) and 33 males (type II). In type I trials, males preferred the female to the male and on average spent 73% (± 7%) of their “mating time” mounting her (Wilcoxon signed rank test with continuity correction, V = 220, *P* = 0.0132). In type 1I trials males preferred the control (i.e. non-extracted) female and on average spent 82% (± 6%) of their “mating time” mounting her (Wilcoxon signed rank test with continuity correction, V = 493, *P* = 0.0002).

### Genetic analysis

We identified 15 isoforms from 13 transcripts in our transcriptome with matches to our query proteins. These transcripts mapped to 16 unique areas of the genome assembly. We blasted these areas against the NCBI nr database (accessed in April 2018) to confirm our annotation. Five of these areas had either no match or matched to an unrelated protein and were thus removed. The remaining genes include 2 putative fatty acid synthases, 2 putative desaturases, 5 putative elongases, 1 putative cytochrome P450 reductase, and 1 putative cytochrome P450-4G1 (Table 3). We found 1042 SNPs within these genes (including 5K upstream and 5K downstream), of these 11 had an F_ST_ value > 0.142 (i.e. in the top 5% of the F_ST_ distribution, Table 4).

**Table 3.** Putative genes involved in CHC synthesis in *C. frigida*

| Scaffold<sup>1</sup> | Gene Position<sup>1</sup> | Best Blast Hit<sup>2</sup> |
| --- | --- | --- |
| scaffold_129 | 185034-197861 | Desaturase 1 |
| scaffold_155 | 109640-112144 | Cytochrome P450-4G1 |
| scaffold_206 | 225893-231312 | Cytochrome P450 reductase |
| scaffold_480 | 98147-106166 | CG5326 (elongase family) |
| scaffold_480 | 2453-12223 | CG31522 (elongase family) |
| scaffold_545 | 1553-3217 | CG5326 (elongase family) |
| scaffold_340 | 1141-16368 | Baldspot (elongase family) |
| scaffold_149 | 7122-16098 | Fatty Acid Synthase 3 |
| scaffold_51 | 250672-259608 | Fatty Acid Synthase 1 |
| scaffold_716 | 36957-55451 | CG2781 (elongase family) |
| scaffold_994 | 14376-17314 | CG9743 (contains Fatty acid desaturase domain and Acyl-CoA desaturase) |

1. From unpublished genome assembly (Wellenreuther *et. al*)
2. From a BLAST against *D. melanogaster* annotated nucleotides on flybase.org

**Table 4.** Outlier SNPs putatively involved in geographic differences

| Scaffold<sup>1</sup> | Position<sup>1</sup> | Variant type<sup>2</sup> | Best Blast Hit<sup>3</sup> | F<sub>ST</sub><sup>4</sup> |
| --- | --- | --- | --- | --- |
| scaffold_129 | 195074 | missense variant | Desaturase 1 | 0.16 |
| scaffold_155 | 116876 | downstream gene variant | Cytochrome P450-4G1 | 0.22 |
| scaffold_206 | 225645 | downstream gene variant | Cytochrome P450 reductase | 0.17 |
| scaffold_206 | 226804 | missense variant | Cytochrome P450 reductase | 0.17 |
| scaffold_206 | 229444 | missense variant | Cytochrome P450 reductase | 0.20 |
| scaffold_206 | 230032 | intron variant | Cytochrome P450 reductase | 0.30 |
| scaffold_480 | 105896 | synonymous variant | CG5326 (elongase family) | 0.22 |
| scaffold_480 | 110918 | downstream gene variant | CG5326 (elongase family) | 0.16 |
| scaffold_994 | 16976 | missense variant | CG9743 (contains Fatty acid desaturase domain and Acyl-CoA desaturase) | 0.16 |
| scaffold_994 | 18844 | downstream gene variant | CG9743 (contains Fatty acid desaturase domain and Acyl-CoA desaturase) | 0.20 |
| scaffold_994 | 19574 | downstream gene variant | CG9743 (contains Fatty acid desaturase domain and Acyl-CoA desaturase) | 0.20 |

1. From unpublished genome assembly (Wellenreuther *et. al*)
2. Annotated in SnpEff (see text for details)
3. From a BLAST against *D. melanogaster* annotated nucleotides on flybase.org
4. Calculated from a separate unpublished data set of WGS from 46 *C. frigida*

## DISCUSSION

Here we explored causes of variation in cuticular hydrocarbons (CHCs) in the seaweed fly *C. frigida*. We detected considerable phenotypic CHC variation attributable to sex, population, and diet and were able to describe genetic variation in candidate genes involved in CHC synthesis that was attributable to population. We show that this trait is likely used as a signal in male mate choice indicating that CHC composition may be under both natural and sexual selection in this system. We discuss these results and their consequences below.

Males and females differed in their CHC composition. Not only did the composition between males and females differ (Table 2, Figure 2-5) but, in addition to that, males consistently had more CHC per gram body weight than females (Figure 4A). Our behavioural trials demonstrate that males preferentially mount females with an intact signal (i.e. CHC cocktail) indicating that CHCs are likely used in male mate choice. Although we were unable to test female choice, it is likely that females also use CHCs in mate choice. A role for CHCs in sexual selection in *C. frigida* would be in line with the general finding that CHCs play a major role in communication in insects, specifically in Diptera (Howard and Blomquist 2005; Blomquist and Bagnères 2010; Ferveur and Cobb 2010). Differences in CHCs have been shown to be involved in a wide variety of behaviours in diptera such as courtship, aggregation, and dominance (Ferveur and Cobb 2010). For instance, 11-*cis*-vaccenyl acetate, a male specific hydrocarbon in *Drosophila melanogaster*, increases male-male aggression while suppressing male mating (Wang and Anderson 2010). The same compound is also involved in aggregation behaviour in both *D. simulans* and *D. melanogaster* (Bartelt et al. 1985; Schaner et al. 1987). Other aggregation hydrocarbons, such as (Z)-10-heneicosene, in *D. virilis* have been shown to attract males and females of certain ages (Bartelt and Jackson 1984). Further behavioural tests in *C. frigida* can determine if CHCs are additionally used in aggregation and male-male interactions.

We found a shift in the composition between sexes, populations, and diets rather than differences in the presence/absence of specific compounds. This type of pattern differs from some dipterans where sex and population differences are mostly qualitative (Carlson and Schlein 1991; Ferveur and Sureau 1996; Dallerac et al. 2000b; Gomes et al. 2008; Everaerts et al. 2010) although many dipterans also show quantitative differences (ex: Jallon and David 1987; Byrne et al. 1995). This discrepancy is likely to be partially explained by the distribution of compounds that make up the CHCs in *C. frigida*. While *C. frigida* chain lengths and compound classes are comparable with other dipteran species (Ferveur and Cobb 2010) the distribution of these compounds is somewhat different. Many dipterans have principal CHC components that make up a large proportion of the CHC composition (Jallon and David 1987; Ferveur and Sureau 1996; Etges and Jackson 2001; Gomes et al. 2008), while the most abundant compound (the C25 alkane) observed in this study only made up on average 15 % of the CHCs in *C. frigida*.

We found strong geographic signatures in CHC composition when flies were raised in a common garden, which indicates a shift in genetic variation. We examined coding variation to determine if we could tie population level phenotypic differences to genetic differences. We observed country level variation with strong differences between Norway and Sweden (Figure 2D, Figure 4, Figure 5). This pattern mirrors a strong genetic split between the two countries (representing approximately 14% of genetic variation, Berdan *et al*, unpublished). We found potential outlier SNPs in desaturases, elongases, a cytochrome P450-4G1, and a cytochrome P450 reductase. Desaturases add double bonds or triple bonds to alkanes (Howard and Blomquist 2005; Blomquist and Bagnères 2010). We found an outlier missense variant in a putative Desaturase 1 as well as a missense and two downstream outlier variants in a putative Desaturase (Table 4). This aligns with our phenotypic data showing differences in the proportion of alkanes vs. alkenes between Norway and Sweden. Elongases lengthen fatty acyl-CoAs and are necessary for chains longer than 16 carbon atoms (Blomquist and Bagnères 2010). We found both synonymous and downstream outlier variants in putative *C. frigida* elongases (Table 4) and corresponding differences in both the mean chain length and variance of the chain length (dispersion around the mean) due to country of origin. P450 reductases are responsible for reducing fatty acids to aldehydes (Blomquist and Bagnères 2010). Knockdown of P450 reductases in *D. melanogaster* leads to a striking reduction of cuticular hydrocarbons (Qiu et al. 2012). P450-4G genes, which are unique to insects, encode an oxidative decarboxylase that catalyzes the aldehyde to hydrocarbon reaction. Work in other insect species has found that a knockdown or knockout of this gene leads to a decrease in overall hydrocarbon levels (Reed et al. 1995; Qiu et al. 2012; Chen et al. 2016). We found a downstream outlier variant in our cytochrome P450-4G1 and multiple kinds of outlier variants in a cytochrome P450 reductase. However, the total amount of CHCs did not differ between populations, only sexes. As males and females share a genome (with the exception of the sex chromosome) it is unlikely that these SNPs affect this difference. Finally, we also found variation between Sweden and Norway in the proportion of methylated compounds. Other studies suggest that differences in the proportion of methylated compounds are often caused by either variation in Fatty Acid Synthase (FAS, de Renobales et al. 1986; Blomquist et al. 1994; Juarez et al. 1996) or multiple copies of FAS with different functions (Chung et al. 2014). We found two different FAS’ in our genome (Table 3) but neither of them contained divergent SNPs. Future studies are needed to examine if these genetic and phenotypic changes are related. Specifically, investigations into both RNA expression and tests of function of these loci will be necessary to provide a direct link. However, the association between the known function of several of these genes (elongases, desaturases) and the corresponding differences in populations are marked. Given that we see shifts in multiple chemical aspects (length, methylation, and alkene/alkane ratio) it is likely that multiple loci are responsible for these patterns.

We also detected plasticity in CHC composition due to the environment; the wrackbed itself had a strong influence on the CHC composition (Figure 3C). Wrackbed composition and microbiome (i.e. the food source for *C. frigida* larvae) varies across *C. frigida* populations in Europe (Berdan, et al., in prep, Day et al. 1983; Butlin and Day 1989; Wellenreuther et al. 2017). Consequently, the CHC composition of *C. frigida* is likely to differ between natural populations in accordance with other research showing that larval diet impacts CHC composition (Liang and Silverman 2000; Rundle et al. 2005; Etges and de Oliveira 2014; Stojkovic et al. 2014).

Together our results indicate that both genetic and environmental factors influence the CHC composition in *C. frigida*. This variation may lead to several potential evolutionary scenarios: 1. Natural selection may select for a different CHC composition in different environments and natural polymorphism in CHC genes may be under direct selection due to this. For example, *Drosophila serrata* and *D. birchii* differ in the environment they inhabit (habitat generalist vs. habitat specialist for humid rainforests). These two species also differ in the concentration of methyl branched CHCs (especially important in desiccation resistance) caused, in part, by changes in the cis-regulatory sequence likely under natural selection (Chung et al. 2014). 2. Natural populations differ strongly in their CHC composition due to environmental effects (i.e plasticity). If assortative mating or selection on preferences is present, this could lead to locally distinct traits and preferences (Chung and Carroll 2015). This would reduce effective migration between populations, as foreign males would be at a disadvantage. Differences in CHC composition and corresponding preferences have been shown to cause behavioral isolation between many different *Drosophila* species or populations (Coyne et al. 1994; Rundle et al. 2005; Etges and de Oliveira 2014).

In conclusion, the analysis of CHC extracts from male and female *C. frigida* from multiple populations revealed a complex and variable mix of more than 100 different compounds. We confirmed that CHCs are likely used in mate choice and found extensive phenotypic variation attributable to diet, sex, and country as well as associated genetic variation attributable to the latter. This reveals that CHC composition is dynamic, strongly affected by the larval environment, and most likely under natural and sexual selection. Further work is needed to explore the evolutionary consequences of these differences.

## Acknowledgements

We thank the members of the Marine Chemical Ecology group at Gothenburg University for helpful comments on experimental design. E.B. was supported by a Marie Skłodowska-Curie fellowship 704920 — ADAPTIVE INVERSIONS — H2020-MSCA-IF-2015. MW and the research were supported by grants from Vetenskapsrådet to MW (2012-3996), from the Mistra Foundation to HP (AquaAgriKelp) and from the Swedish Foundation for Strategic Research to HP (Sweaweed).

