## Supplemental Tables for "Genetic divergence and phenotypic plasticity contribute to variation in cuticular hydrocarbons in the seaweed fly *Coelopa frigida*"

### Supplemental Information

#### **Table S1** –GC-MS instrument settings

| | |
|---|---|
| Injection | 1 µl |
| Inlet | pulsed splitless (1min), 280°C |
| Column | 30m x 0.25 mm I.D. x 0.25µm HP-5MS<br>Max. temperature 325°/350°C<br>Carrier gas: helium<br>Flow: constant flow (0.5 ml/min)<br>Inlet: front inlet<br>Outlet: MSD |
| GC oven | 200°C (3 min), 8°C/min to 325°C, 325°C (10 min) |
| MSD transfer line | 280°C |
| MS scan | 40-600 m/z, 3 min solvent delay |
| MS temperatures | MS Quad 150°C, MS Source 230°C |

**Table S2** – Pair-wise tests for sex and population

| Population differences within females | | | t-Test | | | PERMDISP | |
|---|---|---|---|---|---|---|---|
| | | | t | p-value | | t | p-value |
| Ystad | vs. | Stavder | 1.240 | 0.083 | | 0.939 | 0.462 |
| Ystad | vs. | Skeie | 1.207 | 0.071 | | 0.015 | 0.985 |
| Ystad | vs. | Østhassel | 1.261 | 0.045 | * | 1.232 | 0.321 |
| Stavder | vs. | Skeie | 1.654 | <0.001 | *** | 1.199 | 0.313 |
| Østhassel | vs. | Stavder | 1.565 | 0.002 | ** | 0.467 | 0.708 |
| Østhassel | vs. | Skeie | 0.916 | 0.686 | | 1.528 | 0.212 |

| Population differences within males | | | t-Test | | | PERMDISP | |
|---|---|---|---|---|---|---|---|
| | | | t | p-value | | t | p-value |
| Ystad | vs. | Stavder | 1.919 | <0.001 | *** | 2.316 | 0.085 |
| Ystad | vs. | Skeie | 1.282 | 0.041 | * | 0.102 | 0.935 |
| Ystad | vs. | Østhassel | 1.146 | 0.090 | | 0.658 | 0.594 |
| Stavder | vs. | Skeie | 2.053 | <0.001 | *** | 2.545 | 0.052 |
| Østhassel | vs. | Stavder | 1.855 | <0.001 | *** | 2.070 | 0.087 |
| Østhassel | vs. | Skeie | 1.124 | 0.167 | | 0.640 | 0.624 |

| Sex differences within populations | | | t-Test | | | PERMDISP | | |
|---|---|---|---|---|---|---|---|---|
| | | | t | p-value | | t | p-value | |
| Ystad: F | vs. | M | 1.204 | 0.074 | | 1.039 | 0.368 | |
| Stavder: F | vs. | M | 2.026 | <0.001 | *** | 3.841 | 0.005 | ** |
| Skeie: F | vs. | M | 1.506 | 0.007 | ** | 1.514 | 0.251 | |

**Table S3** – Model significance estimates

| Model | No. predictive components | No. orthogonal components | $R^2X$(cum) | $R^2Y$(cum) | $Q^2$(cum) | RMSEE | p $R^2Y$ | p $Q^2$ |
|---|---|---|---|---|---|---|---|---|
| **Sex and population effects** | | | | | | | | |
| PCA | 8 | | 0.51 | | | | | |
| OPLS-DA for sex | 1 | 3 | 0.36 | 0.76 | 0.57 | 0.25 | 0.001 | 0.001 |
| PLS-DA for population | 4 | | 0.34 | 0.5 | 0.27 | 0.31 | 0.001 | 0.001 |
| OPLS-DA for countries | 1 | 1 | 0.28 | 0.52 | 0.41 | 0.35 | 0.002 | 0.001 |
| **Sex and diet effects in Ystad population** | | | | | | | | |
| PCA | 6 | | 0.54 | | | | | |
| OPLS-DA for sex | 1 | 3 | 0.41 | 0.83 | 0.5 | 0.22 | 0.001 | 0.001 |
| OPLS-DA for wrack type | 1 | 3 | 0.44 | 0.9 | 0.72 | 0.17 | 0.001 | 0.001 |

RMSEE = root mean square error of estimation

**Table S4** – AIC comparison of models for A. Total peak area, B. Proportion of alkenes, C. Proportion of methylated compounds, D. Weighted mean chain length, E. Dispersion around mean chain length.

**A.**

**Model Terms**

| Intercept | Country | Sex | Country*Sex | df | Log Liklihood | AIC | Delta |
|---|---|---|---|---|---|---|---|
| 1.712 | | + | | 3 | -95.545 | 197.3 | 0 |
| 1.657 | + | + | | 4 | -95.117 | 198.6 | 1.31 |
| 1.71 | + | + | + | 5 | -94.716 | 200 | 2.71 |
| 1.915 | | | | 2 | -101.032 | 206.2 | 8.85 |
| 1.86 | + | | | 3 | -100.647 | 207.5 | 10.21 |

**B.**

**Model Terms**

| Intercept | Country | Sex | Country*Sex | df | Log Liklihood | AIC | Delta |
|---|---|---|---|---|---|---|---|
| -1.499 | + | + | | 4 | 170.256 | -332.1 | 0 |
| -1.544 | + | + | + | 5 | 171.11 | -331.6 | 0.5 |
| -1.678 | | + | | 3 | 163.176 | -320.1 | 12 |
| -1.855 | + | | | 3 | 143.044 | -279.8 | 52.26 |
| -2.043 | | | | 2 | 137.998 | -271.9 | 60.23 |

**C.**

**Model Terms**

| Intercept | Country | Sex | Country*Sex | df | Log Liklihood | AIC | Delta |
|---|---|---|---|---|---|---|---|
| -0.2605 | + | + | | 4 | 78.825 | -149.2 | 0 |
| -0.2268 | + | + | + | 5 | 79.104 | -147.6 | 1.65 |
| -0.1558 | | + | | 3 | 76.213 | -146.2 | 3.06 |
| -0.09456 | + | | | 3 | 72.483 | -138.7 | 10.52 |
| 0.008819 | | | | 2 | 70.163 | -136.2 | 13.04 |

**D.**

**Model Terms**

| Intercept | Country | Sex | Country*Sex | df | Log Liklihood | AIC | Delta |
|---|---|---|---|---|---|---|---|
| 26.88 | + | + | | 4 | -121.059 | 250.5 | 0 |
| 27.03 | + | | | 3 | -122.988 | 252.2 | 1.69 |
| 26.92 | + | + | + | 5 | -120.92 | 252.5 | 1.93 |
| 27.16 | | + | | 3 | -127.504 | 261.2 | 10.72 |
| 27.31 | | | | 2 | -129.211 | 262.5 | 12.02 |

**E.**

**Model Terms**

| Intercept | Country | Sex | Country*Sex | df | Log Liklihood | AIC | Delta |
|---|---|---|---|---|---|---|---|
| 8.985 | + | + | | 4 | -209.006 | 426.4 | 0 |
| 8.976 | + | + | + | 5 | -209.005 | 428.6 | 2.21 |
| 8.316 | | + | | 3 | -215.685 | 437.6 | 11.19 |
| 10.16 | + | | | 3 | -227.381 | 461 | 34.59 |
| 9.492 | | | | 2 | -232.159 | 468.4 | 42.02 |

**Table S5** – Type II analysis of deviance tables for best models for: A. Total peak area, B. Proportion of alkenes, C. Proportion of methylated compounds, D. Weighted mean chain length, E. Dispersion around mean chain length.

**A.**

| Term | Df | Chisq | Pr(>Chisq) |
|---|---|---|---|
| Sex | 1 | 11.353 | 0.0007534 |

**B.**

| Term | Df | Chisq | Pr(>Chisq) |
|---|---|---|---|
| Country | 1 | 15.002 | 0.0001074 |
| Sex | 1 | 67.906 | < 2.20E -16 |

**C.**

| Term | Df | Chisq | Pr(>Chisq) |
|---|---|---|---|
| Country | 1 | 5.352 | 0.020699 |
| Sex | 1 | 13.458 | 0.000244 |

**D.**

| Term | Df | Chisq | Pr(>Chisq) |
|---|---|---|---|
| Country | 1 | 13.3253 | 0.0002618 |
| Sex | 1 | 3.8166 | 0.0507463 |

**E.**

| Term | Chisq | Df | Pr(>Chisq) |
|---|---|---|---|
| Country | 1 | 13.841 | 0.0001989 |
| Sex | 1 | 42.808 | 6.04E-11 |
